# Genetic-metabolic coupling for targeted metabolic engineering

**DOI:** 10.1101/156927

**Authors:** Stefano Cardinale, Felipe Gonzalo Tueros, Morten Otto Alexander Sommer

## Abstract

To produce chemicals, microbes typically employ potent biosynthetic enzymes that interact with native metabolism to affect cell fitness as well as chemical production. However, production optimization largely relies on data collected from wild type strains in the absence of metabolic perturbations, thus limiting their relevance to specific process scenarios. Here, we address this issue by coupling cell fitness to the production of thiamine diphosphate in *Escherichia coli* using a synthetic RNA biosensor.

We apply this system to interrogate a library of transposon mutants to elucidate the native gene network influencing both cell fitness and thiamine production. Specifically, we identify uncharacterized effectors of the OxyR-SoxR stress response that limit thiamine biosynthesis via alternative regulation of iron storage and Fe-S-cluster inclusion in enzymes.

Our study represents a new generalizable approach for the reliable high-throughput identification of genetic targets of both known and unknown function that are directly relevant to a specific biosynthetic process.

## INTRODUCTION

A common problem in the field of biotechnology is the unpredictability inherent in engineering complex biological systems. The use of a whole-system approach can prevent some of the obstacles encountered in bioengineering, including toxicity due to metabolite over-production (Fletcher et al., 2016), metabolic bottlenecks (Lechner et al., 2016), and low product titers (Otero and Nielsen, 2013). Systems metabolic engineering relies on the generation and analysis of large datasets to identify key genetic components that contribute to high-yield production phenotypes. However, this methodology often assumes an unperturbed system, and our capacity to elucidate all metabolic and regulatory perturbations affecting the production of a given metabolite is limited.

The impact of engineering a cellular system to produce a primary metabolite can be substantial. The biosynthetic pathways of primary metabolites often utilize specialized enzymes of high catalytic rate and substrate affinity; these attributes support high flux through the pathway, even at low enzyme concentrations (Nam et al., 2012). When these enzymes are over-expressed, the supply of energy, cofactors, and carbon building blocks quickly become limiting and can lead to a strong metabolic burden on the cell. Indeed, metabolic burden can lead to low cell density and reduced titers during the production of recombinant vitamins, such as cobalamin (vitamin B_12_) (Biedendieck et al., 2010) or riboflavin (vitamin B_2_) (Lin et al., 2014). Furthermore, central cell metabolism, which provides building blocks for vitamin biosynthesis, is subject to complex regulation that can affect pathway optimization (Lin et al., 2014).

The effect of producing a metabolite is strongly dependent on the individual metabolite and how it affects cell metabolism and its regulation. Understanding the relevance of this context dependency at the genome level remains a challenge in the field of biotechnology. New approaches enabling high-throughput characterization of fitness and pathway yields are key to achieving this goal. Tagged transposon mutagenesis (Oh et al., 2010) and synthetic RNA-based biosensors (Townshend et al., 2015) are genetic tools that have been applied to study the condition-dependent contributions of genes to cell growth and synthetic pathway output.

Here, we present a methodology for characterizing the metabolic perturbations that are a direct result of the over-production of a primary metabolite in a bacterial cell. Genetic-metabolic coupling is based on the concurrent expression of a synthetic metabolic pathway and an end-product biosensor in a population of bar coded single-gene transposon insertion mutants. Mutant fitness data quantified by deep sequencing are used to identify which genes provide growth advantage exclusively upon selection for the end product. We use genetic-metabolic coupling to assess the influence of *Escherichia coli* cell functions during biosynthesis of a derivative of vitamin B_1_ – thiamine diphosphate (TPP).

We found that the OxyR-Fur-mediated regulation of Fe-S cluster formation strongly affects TPP output. Ultimately, genetic-metabolic coupling allowed the identification of a small set of genes, several of which of uncharacterized or predicted function, with strong population-wide fitness phenotypes from over 2000 tested genes. When introducing specific knockouts in a clean genetic background, the identified genes resulted in the predicted changes to the TPP titers with different production plasmids and in different host strains. We finally demonstrate reverse tuning of several identified genetic targets with multi-copy plasmids further supporting the broad application of genetic-metabolic coupling for metabolic engineering.

## RESULTS

### Coupling population genetics to thiamine metabolic engineering

To create a TPP-overproducing strain for our analysis, we assembled the TPP biosynthetic genes on a plasmid. The genes were arranged in two operons that encoded the two different cellular branches required for the biosynthetic pathway. Four genes are involved in the synthesis of thiazole monophosphate (THZ-P) from L-cysteine, L-tyrosine and deoxy-D-xylulose-5-phosphate (DXP): thiFSGH, *thiC* for the synthesis of 4-amino-5-hydroxymethyl-2-methylpyrimidine (HMP-P) from 5- aminoimidazole riboside (AIR), *thiE* for coupling THZ-P to HMP-PP, and either *thiD*, the kinase that phosphorylates HMP-P to HMP-PP (plasmid pTHId), or *thiM* (plasmid pTHIm), a salvage enzyme that phosphorylates THZ. In the latter case, the strain relies on the native chromosomal *thiD* gene for HMP-P phosphorylation (details are provided in Sup. Fig. 1A-B). The synthesis of TPP relies on several cellular biosynthetic pathways for substrates and in particular purine nucleotides biosynthesis for AIR, cysteine biosynthesis, and S-adenosylmethionine (SAM) and NADPH as cofactors.

To quantitatively relay the intracellular TPP concentration at the single-cell level, we used a TPP biosensor (plasmid pTPP_Bios) based on an engineered TPP riboswitch (Genee et al., 2016). This riboswitch is used to activate the expression of two genes conferring resistance to the antibiotics chloramphenicol (CAM) and spectinomycin (SpeR). This dual selection design allows for the reliable selection of high TPP producers using both the pTHIm and pTHId plasmids with very low false-positive rate (Genee et al., 2016). The extracellular titers of *de novo-produced* TPP in *E. coli* MG1655::ΔtbpAΔthiM (BS134), which carries deletions of the thiamine salvage and import pathways, were ~200 and ~800 μg/l for pTHIm and pTHId, respectively (Fig. 1A). The overexpression of thiamine biosynthetic genes reduced the growth rates by 10% and 35% for pTHIm and pTHId, respectively.

**Figure 1.**
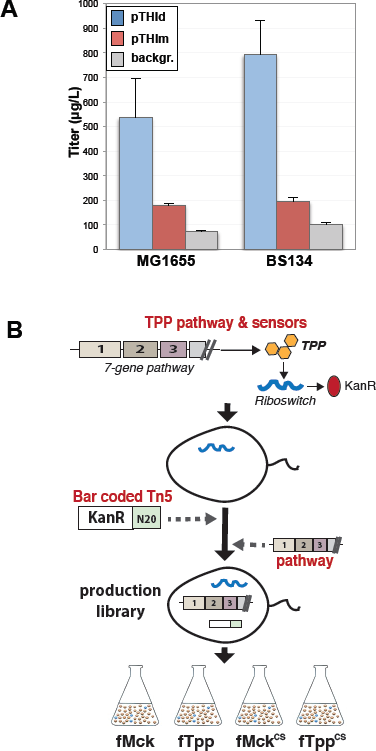
Genetic-metabolic coupling design. **A)** Quantification of extracellular TPP titer with pTHId and pTHIm (cyan and red, respectively) compared to a control (no plasmid, grey) in *E. coli* MG1655 and BS134 strains (n = 4-6) **B)** Graphical scheme of the experimental design. A wild type *E. coli* strain carrying the dual riboswitch (pTPP_Bios) is subject to mutagenesis via a tagged Tn5 transposon. Following library characterization, the TPP production plasmid (pTHIm) or the backbone plasmid (pMck) is electroporated in the pool of mutants to obtain respectively the fTpp and fMck libraries. These libraries are finally used for fitness assays in absence or presence (^CS^) of selection for TPP production.

To understand the cellular components impacting the *E. coli* TPP biosynthetic capacity, we used whole-genome transposon (Tn) mutagenesis to construct libraries of *E. coli* MG1655 Tn5 mutant strains harboring the thiamine selection system (pTPP_Bios) and the plasmid carrying both *de novo* and salvage biosynthetic genes with a lower cell burden (pTHIm), or the plasmid without biosynthetic pathway, constructing the respective libraries lTpp and lMck (Figure 1B). Insertions were evenly distributed across the genome and through the lengths of open reading frames (Sup. Fig. 1C-D) (RefSeq. NC_000913.3). To characterize the impact of transposon insertions within various *E. coli* genes on the intracellular excess of TPP, we subjected the lMck and lTpp mutant libraries to a competitive fitness assay for 16 hours in shake flasks, obtaining robust fitness estimates for 1857 genes in the lMck library and 2054 genes in the lTpp library with high reproducibility across biological replicates (Sup. Fig. 1E). The relative gene fitness in each condition was calculated from strain bar code abundance (log_2_ fitness) using the relative abundance in the Tn libraries at time zero (Wetmore et al., 2015).

### Iron and sulfur significantly impact TPP production

We applied a two-step methodology for identifying genes and biological functions with significant condition-specific fitness phenotype (top/bottom 5%, see also Sup. Methods), or a significantly reduced/increased relative abundance of TPP-producing Tn mutants compared to time zero or mutants carrying the backbone-only plasmid (lMck). First, extreme fitness values were identified by setting top and bottom thresholds for each condition. These values were used to build a factorized matrix with +1 or -1 values corresponding to significantly increased or decreased fitness, respectively, which was later used to establish gene-to-condition associations for functional enrichment (DAVID) (Huang et al., 2008) (Fig. 2A).

**Figure 2.**
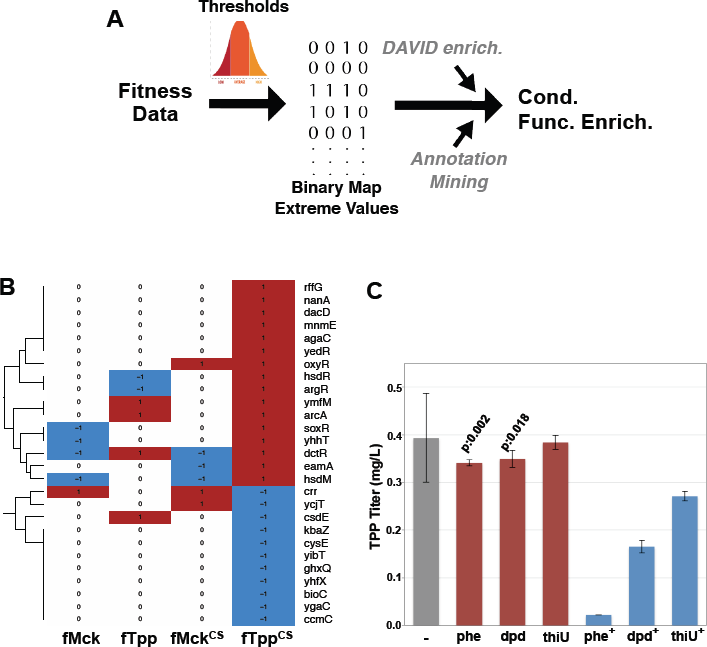
Gene knockouts with significant fitness shift during TPP production. **A)** Data analysis pipeline: significant fitness values are identified and used to build a phenotypic binary map. Functional enrichment in gene clusters is examined by DAVID and by mining RefSeq gene descriptions for functional annotations. **B)** Hierarchical clustering of set of genes with significant fitness shift exclusively during TPP production and selection (right-most column fTpp^CS^, +/- or red/cyan indicate significantly increased/decreased fitness value) (see also Sup. Fig. 2C). **C)** Extracellular TPP concentration from *E. coli* either untreated (-) or grown in presence of low (red bars, phe=20 μM, dpd=30 μM, thiU=100 μM) or high (5x) (cyan bars) concentrations of o-phenanthroline (phe and phe^+^), 2,2’-dipyridyl (dpd and dpd^+^) or thiourea (thiU or thiU^+^) (n = 4-5) (p: two-tailed Student’s t-Test).

We focused our analysis on two effects: the effect of having biosynthetic genes for TPP compared to control backbone-only plasmid and the effect of selecting for intracellular TPP compared to no selection. In both cases we expected that the deletion of a gene encoding factors important for the biosynthesis of TPP, which was not provided in the cultivation media, or for replenishing cellular resources would lead to relatively reduced fitness compared to the full population.

In the presence of the TPP heterologous pathway but without selective pressure (fTpp), we identified 138 genes with significant fitness changes, of which 102 exhibited reduced and 36 exhibited increased fitness (Sup. Fig. 2A and Sup. Table 1). Knockouts of genes affecting cofactor or substrate supply to the TPP pathway showed fitness defects: *rtn* (ΔF: -1.0, ΔF: fitness difference to control) and *fucA* (ΔF: -1.3), implicated in ribose metabolism, *bfr* (ΔF: -1.1) and *yaaA* (ΔF: -0.8) in iron homeostasis, and *dxs*, involved in DXP biosynthesis. We also found enrichment for knockouts of genes involved in protein folding (Sup. Table 1, *P* < 0.05, with Bonferroni correction for multi-hypotheses testing). During selection for TPP without the TPP heterologous pathway (fMck^CS^), several additional genes involved in cellular iron homeostasis (*fes, ftnA* and *cueO*) and the Gene Ontology (GO) class “water-soluble vitamin biosynthesis functions” (GO:0006767, *P* < 0.05, Sup. Fig. 2B and Sup. Table 1) showed reduced fitness. Overall, this result indicates that TPP biosynthesis relies strongly on the cell protein and vitamin synthesis apparatus, especially cellular iron availability. This result also highlights that our synthetic TPP biosensor is sensitive to the concentration of endogenous TPP.

We next studied the effect of selecting for TPP in the presence of the heterologous TPP pathway (fTpp^CS^). We compiled a list of 45 genes with significant fitness changes, of which 27 were specific to this condition (e.g., with a neutral phenotype in the absence of selection or biosynthetic genes; Fig. 2B and Sup. Fig. 2C-D). This set of genes was functionally enriched for iron and glutathione ABC transporters (Kyoto Encyclopedia of Genes and Genomes [KEGG] pathway, DAVID Bonferroni-corrected *P* < 0.01), involvement in the sulfur-compound biosynthetic process (GO:0044272 *P ≤* 0.05), and DNA restriction and modification (GO:0009307, *P* < 0.01). In addition, similar phenotypes were shared by functionally related genes: reduced Fe-S cluster availability (*csdE, cysE, ccmC* mutations) presented reduced fitness (Fig. 2B –bottom), while gene knockouts that were involved in sulfur assimilation, major redox regulators OxyR and SoxR, and HSD DNA restriction enzymes resulted in a positive fitness shift (Fig. 2B). Indeed, the pTHId pathway was more stable in a ΔhsdMR knockout compared to wild type (Sup. Fig. 3), possibly suggesting direct cleavage by this endonuclease.

SAM turnover during TPP biosynthesis requires extensive NADPH reductive power (Chen et al., 2015). NADP+ can increase cellular oxidation, impair Fe-S cluster formation, and cause accumulation of hydrogen peroxide and reactive oxygen species (ROS) all tightly controlled by OxyR-SoxR regulons (Imlay, 2015). We tested the effect on TPP titer from limiting Fe-S clusters and the activation of OxyR-SoxR regulons. Sequestration of intracellular unbound iron by iron chelators o-phenanthroline and 2,2’- dipyridyl (Chupka et al., 1988) led to a small but significant reduction in TPP titer already at low concentrations (~25 μM) whereas thiourea showed near-normal TPP levels (Fig. 2C, P < 0.02, red bars). These effects were more significant at 5-fold higher compounds concentration, (Fig. 2C, cyan bars), indicating that free iron and Fe-S clusters are limiting during TPP production and the removal of ROS has an effect similar to the OxyR knockouts and regulon down-regulation.

### Elucidation of the Fe-S cluster biogenesis network

We used a machine-learning algorithm to mine RefSeq gene descriptions (Tatusova et al., 2014) and quantified for each condition the effects of knocking out genes with biological functions related to iron, sulfur, cysteine, tyrosine and ribose, which are cofactors or substrates involved in TPP biosynthesis (Chatterjee et al., 2008; Kriek et al., 2007). We also looked at cellular redox regulation, which is believed to influence cellular vitamin metabolism (Dougherty and Downs, 2006). We identified 108 genes in the *E. coli* genome related to these functions, among which 57 genes (~53%) showed significant fitness shifts during selection for TPP production: 35 were associated with cofactor supply, and 25 were associated with cellular redox regulation (fTpp^CS^ and fTPP, *P* < *0.1* compared to a random set by random sampling with replacement or bootstrapping, Fig. 3A). The presence of a significant number of genes involved in redox regulation suggests that oxidative stress can be relevant during TPP production.

**Figure 3.**
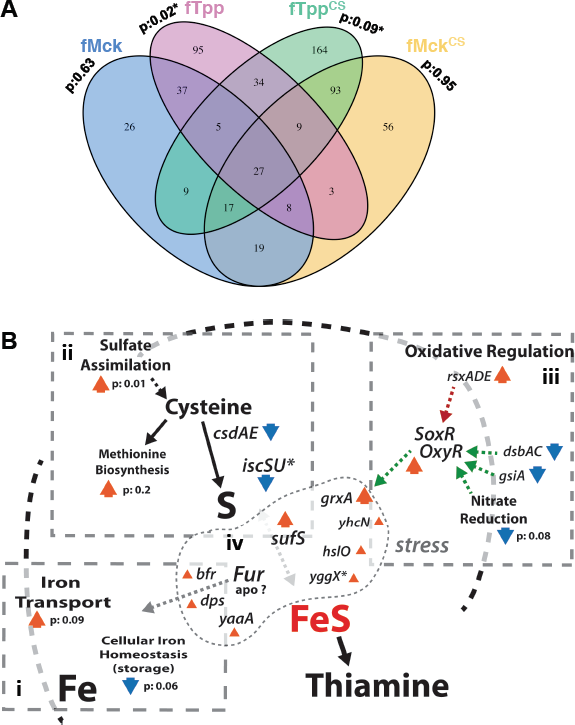
Diverse effects of Fe-S clusters metabolism. **A)** Venn diagram of sets of genes with significant fitness for each cultivation condition (p: the P-value of Fe-S metabolism enrichment for each condition found via mining gene annotations, *: significant, bootstrap n = 10000; null hypothesis: enrichments can occur with the same probability in a random set of genes of equal size). **B)** A map of the Fe-S biogenesis and regulatory network. Four sub-networks are represented: iron supply (*i*), sulfur supply (*ii*), redox regulation of oxyR-soxR regulons (*iii*) and Fe-S metabolism (*iv*) (orange/cyan arrowheads: improved/reduced fitness, small-size: 1-2 St. Dev. from mean, big-size: > 2 St. Dev. from mean) (see also Sup. Methods and Table 2).

To extend our analysis beyond a specific enzyme or transporter, we quantified the roles of the production phenotypes of parent GO classes (Keseler et al., 2009) associated with genes with extreme fitness phenotypes. We found that mutants of ‘Cellular Iron Ion Homeostasis’ (GO: 006879) genes had a significant low-fitness phenotype during selection for TPP (*P* < 0.05, Sup. Table 2); however, mutations in genes involved in iron, siderophores and ferric-enterobactin transport (GO: 00015682, 0015684, 0015685, 015891) were associated with significantly higher fitness (P < 0.1, Fig. 3Bi and Sup. Fig. 4A-B).

In bacteria, sulfur is mainly assimilated in the cell in a reduced form, as hydrogen sulfide from sulfate, and is incorporated into the amino acids cysteine and methionine. We found that the average fitness of mutants in genes involved in ‘Sulfate Assimilation’ (GO: 0000103) had significantly increased (P < 0.01) during selection for TPP (Fig. 3Bii, Sup. Fig. 4C). We also found that impairment of genes related to ‘Methionine Biosynthetic Process’ (GO:0009086) but not cysteine biosynthesis (ΔcysE: -1.5 ΔF) led to increased fitness (P < 0.1) (Sup. Fig. 4D-E). In *E. coli*, the sulfur found in Fe-S clusters is derived from cytosolic L-cysteine mainly through the enzyme *IscS* (cysteine desulfurase) and the conversion of L-cysteine to L-alanine. We could not obtain fitness data for members of the ISC complex, likely because of its essentiality, but the deletion of the second sulfur-accepting complex – the CSD sulfur transfer system – showed reduced fitness upon selection for TPP (ΔF, Δ*csdA:* -0.9, Δ*csdE:* - 3.4) (Fig. 3Bii). Surprisingly, we found that mutants in the SUF system for iron-sulfur cluster assembly that is employed under iron starvation or oxidative stress conditions (Outten et al., 2004) exhibited higher fitness (ΔF, Δ*sufS:* +0.4, Δ*sufA:* +0.7) (Fig. 3Bii).

The process of inclusion of Fe-S clusters in enzymes in *E. coli* is highly sensitive to the environment and its redox state (Crack et al., 2014; Ding et al., 2005). Transposon insertions in *soxR* and *oxyR* both showed higher fitness during selection for TPP (ΔF, respectively, +1.0 and +1.13). In addition to the *suf* operon, we found that insertions in other components of this response pathway had similar effects: the iron chelators *yaaA* and *dps* (ΔF: +1.4 and +0.6, respectively), *yhcN* (ΔF: +0.6), which is induced during peroxide stress (Lee et al., 2010), and the chaperone Hsp33 (ΔF: Δ*hslO* +0.4). During oxidative stress, both OxyR and SoxR upregulate the *fur* regulon that is regarded to play a key role in maintaining the stability of cellular iron levels (Seo et al., 2014). Ferritin and enterobactin have respectively key iron storage and iron uptake functions in *E. coli.* Bacterioferritin, whose expression is controlled by Fur via the small regulatory RNA RyhB, is important for sequestering iron thus limiting cellular oxidative damage caused by Fenton chemistry (Bou-Abdallah et al., 2002). We found that while Tn insertions in iron-storage ferritin and enterochelin esterase, which releases iron from ferric enterobactin, resulted in fitness defects (ΔF: Δ*ftnAB* ~ -2.0, Δ*fes* -3.3), the lack of bacterioferritin had a positive effect on fitness during selection for TPP (ΔF: Δ*bfr* +1.0) (Fig. 3Biv).

We found that the control of OxyR-SoxR activation also had a significant effect. OxyR is activated by oxidation, likely by oxidized glutathione, for which just a Cys^199^ – Cys^208^ disulfide bond appears to be sufficient (Georgiou, 2002). We found that the impairment of glutathione import to the cytosol and *E. coli* disulfide bond system enzymes all result in negative fitness (ΔF, Δ*gsiA*: -1.4, Δ*dsbA*: -2.0, Δ*dsbC*: -1.7). In contrast, the lack of *soxR* inactivation by the RSX Membrane Reducing System (Koo et al., 2003) showed positive growth phenotypes (ΔF, Δ*rsxA*: +1, Δ*rsxD*: +0.8, Δ*rsxE*: +0.6) (Fig. 3Biii).

Overall, our data suggest that TPP overproduction likely causes oxidative stress as previously reported (Kriek et al., 2007) and probably iron deprivation, resulting in an activation of the OxyR-SoxR and possibly Fur regulons, which could be responsible for the distinct effects on iron uptake and sequestration (Fig. 3Biv and i) due to negative feedback (Seo et al., 2014).

### Population dynamics effectively identify metabolic targets

We set out to examine whether population-based screening could be used to reliably identify metabolic targets by coupling genetic modification to metabolism. We measured the extracellular titers of TPP produced *de novo* from the plasmid pTHId in 24 genomic deletions of *E. coli* MG1655 BS134 strains selected from the characterized Fe-S biogenesis network (Fig. 3B). For comparison with the newly identified genetic targets we selected the following five loci previously shown to affect thiamine biosynthesis: *mrp* (Boyd et al., 2008), *purF* (Frodyma et al., 2000), *iscAU* (Agar et al., 2000), *iscR* (Giel et al., 2013) and *yggX* (Gralnick and Downs, 2003). We expected to see reductions in biosynthetic output for Δ*mrp*, Δ*purF* and Δ*iscAUS*, and an increase for Δ*iscR*, a regulator that inhibits transcription from the *iscAUS* operon (Giel et al., 2013). We created 14 single-gene knockouts and 10 multi-gene locus lesions (Sup. Table 3). For the latter, we expected to observe an additive effect of the genes on TPP output because their products did not appear to be part of enzyme cascades (Sup. Table 4).

We observed strong correlation between the shift in fitness of a gene knockout in the multiplexed competitive growth assays and the extracellular TPP titer of the isolated gene knockout (correlation: 0.89 excluding the five test loci). Iron supply and availability strongly contributed to TPP output. Disruption of iron storage in ferritin and mobilization from enterobactin completely disrupted the TPP titer (Δ*ftnB* and Δ*fes-entF*, Fig. 4A). In contrast, knockouts of two genes induced by Fur-OxyR regulons – *bfr* and *yaaA* – whose mutants showed fitness improvements in competitive assays (Fig. 3Bi), yielded higher extracellular TPP titers (Δ*bfr*: +0.3 ±0.2, Δ*yaaA*: +0.4 ±0.1, fold difference from mean titer, see also Sup. Fig. 5) compared to the average titer across all knockouts and wild type.

**Figure 4.**
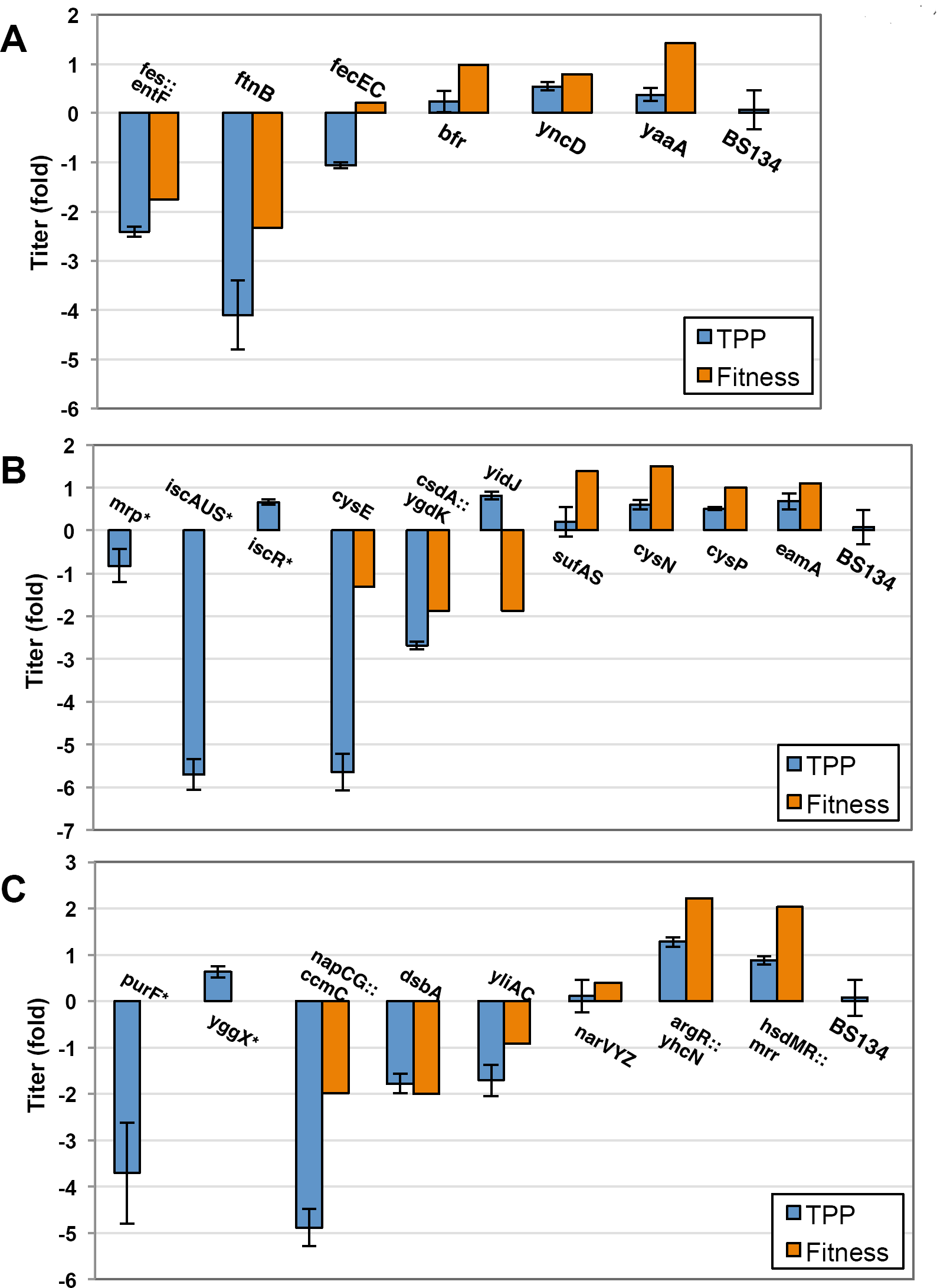
TPP quantification in a set of genetic targets. Extracellular titers for individual locus deletions compared to the average titers across all strains (fold difference, for absolute values see Sup. Fig. 5). Strains are grouped in three different sub-networks involved in Fe-S biogenesis: iron supply (**A**), sulfur supply (**B**) and oxidative regulation (**C**). (*: Test KO, fitness error bars are reported for multi-gene loci) (Error bars: St. Dev. n = 3-5).

We then investigated pathways involved in S-incorporation into Fe-S in *E. coli*, including free cytosolic cysteine. TPP output decreased as expected from previous reports (Agar et al., 2000; Boyd et al., 2008; Giel et al., 2013) in the two test loci Δ*mrp* and Δ*iscAUS* (respectively, -0.8 ±0.04 and undetected, fold difference). In contrast, knockout of the repressor *iscR* resulted in a higher TPP titer (+0.7 ±0.06, Fig. 4B). We found that disrupting cysteine formation and the mobilization of its sulfur by the cellular sulfur transfer system CSD significantly decreased the TPP titer compared to the mean knockout effect (Δ*cysE:* undetected, Δ*csdA-ygdK:* -2.7 ±0.09), which is in agreement with the reduced fitness of mutants in its components (Fig. 3Bii). We previously found that mutants in the *E. coli* SUF sulfur transfer system and in various proteins involved in sulfate assimilation and export exhibited improved growth during TPP selection (Fig. 3Bii). The TPP output of single-locus knockouts agreed with these observations: the average extracellular TPP titers were higher in these knockouts compared to other knockouts and wild type (Δ*sufAS*: +0.4 ±0.2, Δ*cysN*: +0.6 ±0.1, Δ*cysP*: +0.5 ±0.04, Δ*eamA*: +0.7 ±0.2, fold difference) (Fig. 4B).

The remaining tested loci encoded proteins involved in redox control of a number of cellular functions, particularly iron homeostasis, which is tightly regulated by the SoxR-OxyR-Fur regulons (Jang and Imlay, 2011; Seo et al., 2014). The OxyR regulon is activated by an environment that favors disulfide bonds, in which glutathione plays a key role (Georgiou, 2002), and by nitric oxide via S-nitrosylation (Seth et al., 2012). Deletion of the narVYZ locus, for which a mild fitness effect was calculated (ΔF: +0.4), showed TPP titers close to the knockouts average (Δ*narVYZ;* 0.10 ±0.3) (Fig. 4C). In contrast, we found that knockouts of the three predicted activators of the OxyR regulon exhibited significantly reduced TPP titers: Δ*dsbA*, which is involved in disulfide bond formation (Depuydt et al., 2007) (-1.8 ±0.2), Δ*yliAC* (-1.7 ±0.3) and Δ*napCG* (undetected). Mutants in these loci showed reduced fitness during TPP production, whereas those in the SoxR-OxyR regulons had higher fitness. These observations suggest a role for the SoxR-OxyR regulon in diminishing thiamine biosynthesis. Indeed, knockouts of three downstream genes activated by OxyR - the peroxide-response genes *yhcN* (Lee et al., 2010) and *yggX* (Gralnick and Downs, 2003) - had higher extracellular TPP titers (ΔyggX: +0.6 ±0.1, Δ*argR-yhcN*: +1.3 ±0.1, fold difference) (Fig. 4C), in agreement with the shifts in fitness of their pooled mutants (Fig. 3Biii and Sup. Table 3).

### Tuning of central Fe-S control

We examined the role of specific regulators of the complex *E. coli* SoxR-OxyR-Fur regulons during TPP production. We quantified the extracellular TPP titer obtained with Δ*soxR, ΔoxyR* and Δ*fur* single-gene knockouts introduced in *E. coli* strain TOP10, which lacks DNA restriction systems, and the production strain BS134. We found that compared to the background strains, deletion of SoxR did not lead to increased TPP titers, but the deletion of both OxyR and Fur dramatically increased pathway output in both TOP10 and BS134 (Figure 5A-B, respectively). This result pinpointed the OxyR-Fur regulon, which directs iron homeostasis (Seo et al., 2014), as central cellular response to TPP production.

**Figure 5.**
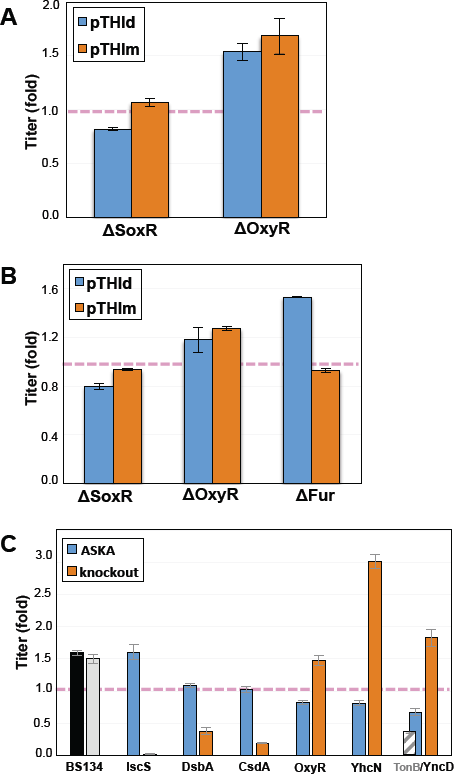
Reverse-engineering OxyR and Fe-S regulation. Extracellular titers obtained by engineering central Fe-S cluster formation and the cellular stress response. (**A**-**B**) TPP titer measured in knockouts of central oxidative stress response in *E. coli* TOP10 (A) and BS134 (B) strains bearing pTHIm and pTHId production plasmids (respectively cyan and orange, pink dashed line: background strain) (n = 6-8). (**C**). Extracellular TPP titer quantified in *E. coli*BS134 over-expressing several identified genetic targets in Fe-S cluster metabolism and oxidative response (cyan bars) compared to their deletion (knockouts, orange bars) (Pink dashed line: average production) (n = 6-8).

In order to determine whether this response and Fe-S cluster formation could be tuned via the identified genetic targets, we over-expressed the following genes as GFP-fusions (Kitagawa et al., 2005): *iscS, dsbA, csdA, oxyR, yhcN, yncD* and *tonB*, in the same strain BS134 used for knockouts. Compared to knockouts, TPP titers resulting from gene over-expression showed an opposite shift. In particular, the titer increased during over-expression of *iscS* and *dsbA* while decreased during over-expression of *oxyR, yhcN* and *yncD* (Figure 5C).

## DISCUSSION

The study shows that by coupling the growth of genome-wide bar coded transposon mutants to a biosynthetic product it is possible to identify product-specific metabolic and regulatory bottlenecks. We show that disruptions in the assimilation of certain forms of iron and sulfur led unexpectedly to improved fitness upon selection for TPP production (Fig. 3Bi and 3Bii), a result that was supported by measuring the extracellular TPP titer from individual gene knockouts (Fig. 4). Further, over-expression of *tonB*, involved in the uptake of iron-siderophore chelates, led to a significant decrease in TPP titer (Fig. 5C). We included in our validation genes that are poorly characterized or of only predicted function, which constitute the so-called Y-genes within the *E. coli* genome, and have strong fitness phenotypes with regard to TPP production. Two genes in particular have predicted function in siderophore uptake *(yncD*, a TonB-dependent receptor) and aryl-sulfate assimilation (*yidJ*, a predicted sulfatase). Both *yncD* and *yidJ* knockouts exhibited TPP titers 20-25% higher than that of wild type cells (Fig. 4), further confirming that certain assimilatory activities may play a role in inhibiting TPP biosynthesis possibly through the regulator Fur.

Applied to TPP biosynthesis, genetic-metabolic coupling uncovered that the regulation of Fe-S homeostasis mediated by OxyR-Fur regulons diminishes biosynthetic output by depleting or sequestering cellular Fe-S clusters. In fact, Fe-S clusters are critical for the function of TPP pathway enzymes ThiC and ThiH. The effects of changing global regulators are notoriously idiosyncratic. Nevertheless, TPP-coupled fitness data of mutants in the regulons could be reproduced in different *E. coli* strains and production plasmids (Fig. 5A-B). Although mutants were not screened for fitness against wild type cells at the population level, 9 of 10 selected high-fitness mutants when introduced in wild type exhibited an extracellular TPP titer 0.2- to 2-fold higher (with the exception of narVYZ), while excluding yidJ all other low-fitness mutants exhibited significantly lower titers (Fig. 4). We also demonstrate that population-wide mutant fitness data could serve as basis for tuning a heterologous pathway through over-expression of genetic targets from multi-copy plasmids (Fig. 5C).

The characterization of how important parts of the *E. coli* stress response and nutrient assimilation affect a complex heterologous pathway and the identification of poorly characterized genes, which today still represent ~30% of the *E. coli* genome, contributing to this response demonstrate the power of genetic-metabolic coupling for the identification of target genes for the metabolic engineering of a specific biosynthetic product.

## EXPERIMENTAL PROCEDURES

### Strains, plasmids and cultivation media

Plasmid DNA, PCR products and gel extractions were prepared with appropriate kits (Qiagen ©). Synthetic oligonucleotides were purchased from Integrated DNA Technologies (IDT, Inc). In-frame GFP fusions of selected genes for expression from multi-copy plasmid were obtained from the ASKA collection (Kitagawa et al., 2005). Single-locus deletions were constructed by employing lambda Red recombineering. The thiamine selection system (pTPP_Bios) was described in a previous study from our laboratory (Genee et al., 2016). The TPP biosynthetic pathway was assembled from native *E. coli* MG1655 genes via PCR amplification. A detailed description of the molecular tools is provided in Sup. Info.

### Analytical quantification of TPP

Extracellular TPP concentration was measured using a modified thiochrome-high-performance liquid chromatography (HPLC) assay procedure. Briefly, fresh colonies carrying either pTHId or pTHIm were inoculated into 1 mL MOPS rich media lacking thiamine (see above) and containing 30 μg/mL chloramphenicol and 50 μg/mL spectinomycin in deep-well culture plates and then cultivated twice to saturation in a 48-hour period (Sup. Info.).

### Construction of Tn-mutants, TnSeq and BarSeq protocols

For construction of tagged transposon mutants of *E. coli* MG1655 and bioinformatic analysis of NGS data we relied on reported methodology and tools (Wetmore et al., 2015). BarSeq reads are converted into a table reporting the number of times each bar code is observed using a custom Perl script (MultiCodes.pl). Given a table of bar codes, where they map in the genome, and their counts in each sample, strain and gene fitness values are estimated with a custom R script (FEBA.R) (Sup. Info.). The bioinformatic tools can be accessed at https://bitbucket.org/berkeleylab/feba.

### Calculation of gene fitness

The gene fitness is the weighted average of Tn mutant strains fitness within the gene, that is strains with more reads have less noisy fitness estimates and are weighted more highly. In more detail, we first selected a subset of strains and genes that have adequate coverage in the time-zero samples (2 reads per strain and 20 reads per gene are considered adequate). Only strains that lie within the central 10 to 90% of a gene coding region are considered. Then, for each sample:

Strain fitness = 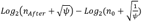

Where Ψ is a “pseudocount” to control for very noisy estimates when read counts are very low. (Further details can be found in Sup. Info. and Wetmore et al., 2015).

## ABBREVIATIONS

KO: knockout, Tn: transposon, aa: amino acid, DNA: deoxyribonucleic acid, RNA: ribonucleic acid, RBS: ribosome binding site.

## SUPPLEMENTAL INFORMATION

Refer to the web version on PubMed Central for supplementary materials.

## AUTHOR CONTRIBUTIONS

Conceptualization, S.C. and M.O.A.S.; Methodology, S.C. and M.O.A.S.; Investigation, S.C. and F.G.T.; Writing - Original Draft, S.C.; Writing - Review & Editing, S.C. and M.O.A.S.; Funding Acquisition, M.O.A.S.

## Competing financial interests

The authors have no competing financial interests to declare.

## ACKNOWLEDGMENTS

This research was funded by the European Union Seventh Framework Programme (FP7-KBBE-2013-7-single-stage) under grant no. 613745 “Promys” and the Novo Nordisk Foundation (NNF) grant no. 11355-444 “Biobase”. F.G.T acknowledges NNF Ph. D. fellowship grant no. NNF16CC0020908. M.O.A.S acknowledges additional funding from the NNF.

We thank Dr. Adam Deutschbauer (Lawrence Berkeley National Laboratory - Berkeley, USA) for kindly providing strains and plasmids utilized here for transposon mutagenesis. We also thank H. Genee and L. Gronenberg (Biosyntia ApS) for donating the source molecular parts for producing and selecting thiamine derivatives and for establishing analytical protocols. We also thank Dr. Lei Yang and Pernille Smith (iLoop - DTU Biosustain) for constructing strain BS134, and Dr. Anna Koza for technical support with Illumina sequencing.

